# SumVg: Total heritability explained by all variants in genome-wide association studies based on summary statistics with standard error estimates

**DOI:** 10.1101/016857

**Authors:** Hon-Cheong So, Pak C. Sham

## Abstract

Genome-wide association studies (GWAS) have become increasingly popular these days and one of the key questions is how much heritability could be explained by all variants in GWAS. We have previously proposed an approach to answer this question, based on recovering the “*true*” *z-*statistics from a set of *observed z*-statistics. Only summary statistics are required. However, methods for standard error (SE) estimation are not available yet, thereby limiting the interpretation of the results. In this study we developed resampling-based approaches to estimate the SE and the methods are implemented in an R package. We found that delete-*d*-jackknife and parametric bootstrap approaches provide good estimates of the SE. Methods to compute the sum of heritability explained and the corresponding SE are implemented in the R package SumVg, available at https://sites.google.com/site/honcheongso/software/var-totalvg

## 1. INTRODUCTION

Genome-wide association studies (GWAS) have proven to be successful in dissecting the genetic basis of a variety of diseases. A number of new susceptibility loci have been discovered, providing novel insight into the pathophysiology of many diseases. Nevertheless, a large proportion of the heritability still remained unexplained. It is natural to question the maximum variance that could be explained by all variants in a GWAS (or meta-analyses of GWAS), as we expect many true susceptibility variants are “hidden” due to limited power.

Yang et al (2010) derived a method to estimate the variance explained by all SNPs in a GWAS by a liner mixed model with random SNP effects. We have developed an alternative framework to achieve the same goal requiring only the summary statistics. Essentially, we aimed to recover the “*true*” *z-*statistic from a set of *observed z*-statistics based the following formula established by Brown (1971) and Efron (2009). The corrected *z*-statistics are then converted to variance explained. This approach does not rely on any distributional assumptions of the effect sizes of susceptibility variants. Our method has been applied in a number of studies [for example see (Benke, et al., 2014; Lubke, et al., 2012; van Beek, et al., 2014)]. As we have discussed in our previous work (So et al., 2011), if raw data is available, a standard non-parametric bootstrap (i.e. sampling individuals with replacement) could be employed to estimate the standard error (SE). However, in many cases only summary statistics are available and there are currently no methods for evaluating the SE of the total heritability explained.

In this paper we proposed several resampling approaches to estimate the SE of the total heritability by all SNPs in GWAS, based on summary statistics. The methods are implemented in the R package SumVg.

## 2. METHODS

### 2.1 Estimation of the total variance explained

Readers may refer to our previous paper (So, et al., 2011) for details on estimation of the sum of heritability explained. In brief, we estimated the “true” *z*-statistics by the following correction formula:

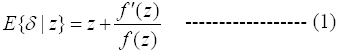

where *z* denotes the observed *z*-statistic and *δ* denotes the “true” z-statistic (i.e. the *z*-statistic one would obtain if there were no random noise; it reflects the actual effect size).

We also proposed previously an alternative approach by evaluating the expected effect size conditioned on H_1_. The “true” z-statistic is estimated by dividing the estimator (1) by [1- *fdr*(*z*)], where fdr is the local false discovery rate described in Efron (2001).

The conditional estimator is however prone to relatively large random variations as it involves local fdr estimation of each SNP. In subsequent applications of our heritability estimation method (Benke, et al., 2014; Lubke, et al., 2012; van Beek, et al., 2014), the unconditional estimator (1) was employed. We shall hence focus on the unconditional estimator in this paper, although the resampling approaches described below can readily be applied to other estimators in our previous work (So, et al., 2011) as well.

### 2.2 Standard and delete-d-jackknife

In a standard jackknife procedure (Miller, 1974), we estimate the standard error (SE) by leaving out one observation at a time. The SE is defined by

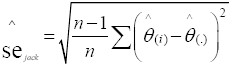

where *n* is the sample size, 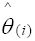 is the parameter estimate from the sample with the *i* th observation removed and

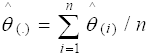

In our case the parameter is the sum of heritability from all variants.

An extension is the delete-*d*-jackknife (Shao and Wu, 1989) where we leave out *d* observations at a time. There are in total *N*=_*n*_C_*d*_ possibilities of removing *d* out of *n* observations. In practice, *N* is usually very large. One may simply randomly repeat the procedure *m* times only (*m* ≤ *N*) instead of exhausting all possibilities of removing *d* out of *n* observations.

The standard error is given by

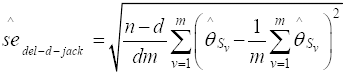

There is no consensus on the best choice of *d*. Chatterjee (1998) suggested *n*/5 as a reasonable choice for *d* based on consideration of efficiency and likely model conditions. We followed the suggestion by Chatterjee (1998) and set *d* as *n*/5 (=20000) in our simulations.

### 2.3 Bootstrap approaches

#### 2.3.1 Non-parametric bootstrap by resampling summary test statistics

SE was estimated by sampling the *z*-statistics with replacement (Efron, 1979). A similar strategy of resampling summary statistics has been employed previously in Storey (2002), but it was used for estimating the SE of false discovery rates.

#### 2.3.2 Parametric bootstrap

We proposed three methods to estimate the SE based on a parametric bootstrap approach. In the first method, in each replication we simulated *z*-statistics based on 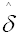, the corrected *z*-statistics from original sample. We have

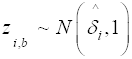

where *z*_*i,b*_ denotes the *i* th *z*-statistic in the *b* th bootstrap replicate.

We further proposed a modified approach by also considering the local fdr of each *z*-statistic. In each replicate, we simulate *z-*statistics according to the following scheme:

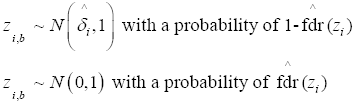

Alternatively, one may employ the original *z*-statistics instead of the corrected *z*-statistics as the mean in each simulation, i.e.

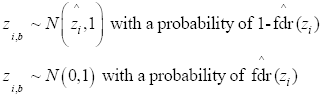

The standard error is then computed from the simulated *z*-statistics.

### 2.4 Tests of resampling-based SE estimates

We compare the SE estimated from the above methods with the “true” SE obtained from two hundred simulations with known data generating distributions. The details of the simulations were described in our previous paper (So, et al., 2011).

Two hundred replicates were run for each bootstrap or jackknife procedure. We focus on quantitative traits in our simulations but the results should apply to binary traits as well, as the only difference in these two scenarios is the formula to convert *z* to variance explained (Vg).

## RESULTS AND DISCUSSIONS

The results are shown in table 1. The standard non-parametric bootstrap approach performed the worst among all methods, producing inflated estimates of SE. The standard (delete-1) jackknife worked reasonably well when the total heritability explained is high (when heritability = 0.295), but tends to overestimate the SE when the total heritability is lower. The delete-[*n*/5]-jackknife on the other hand performs better at all levels of heritability. This may be explained by the fact that the sum of Vg is not a very smooth parameter. The other methods including parametric bootstrap and the modified versions with consideration of local fdr performed reasonably well and closely resemble the true parameter estimates.

**Table 1.** Standard error (SE) of the sum of variance explained estimated by different resampling approaches

| Sum of Vg | Sample size | Mean | True SE | Jack del-1 | Jack del-($n/5$) | Para boot | Wt boot1 | Wt boot2 | Non-para boot |
| --- | --- | --- | --- | --- | --- | --- | --- | --- | --- |
| 0.101 | 5000 | 0.196 | 0.0481 | 0.0638 | 0.0535 | 0.0476 | 0.0544 | 0.0450 | 0.0875 |
|  | 10000 | 0.117 | 0.0254 | 0.0323 | 0.0237 | 0.0227 | 0.0281 | 0.0218 | 0.0364 |
|  | 20000 | 0.098 | 0.0146 | 0.0095 | 0.0131 | 0.0154 | 0.0155 | 0.0155 | 0.0198 |
| 0.191 | 5000 | 0.206 | 0.0495 | 0.0949 | 0.0583 | 0.0531 | 0.0504 | 0.0511 | 0.1039 |
|  | 10000 | 0.148 | 0.0260 | 0.0473 | 0.0334 | 0.0252 | 0.0266 | 0.0252 | 0.0547 |
|  | 20000 | 0.158 | 0.0151 | 0.0473 | 0.0155 | 0.0166 | 0.0141 | 0.0148 | 0.0235 |
| 0.295 | 5000 | 0.231 | 0.0487 | 0.0538 | 0.0428 | 0.0538 | 0.0484 | 0.0476 | 0.0685 |
|  | 10000 | 0.213 | 0.0269 | 0.0279 | 0.0310 | 0.0333 | 0.0294 | 0.0322 | 0.0450 |
|  | 20000 | 0.244 | 0.0146 | 0.0155 | 0.0190 | 0.0158 | 0.0159 | 0.0153 | 0.0292 |

Vg, variance explained;
jack del-1, delete-1-jackknife;
jack del-( $n/5$ ), delete- $d$ -jackknife with $d$ equal to sample size divided by 5;
para boot, parametric bootstrap approach as described in the text;
wt boot1, a "weighted" bootstrap approach with consideration of the local $\text{fdr}$ , using the *observed* $z$ -statistic as the mean in each simulation;
wt boot2, a "weighted" bootstrap approach with consideration of the local $\text{fdr}$ , using the *corrected* $z$ -statistic as the mean in each simulation;
non-para boot, non-parametric bootstrap by sampling the $z$ -statistics with replacement.

In conclusion, we have proposed several resampling approaches to derive the SE of the total heritability explained in GWAS. The delete-[*n*/5]-jackknife and parametric bootstrap methods provided reasonably good estimates of SE.

It should be noted that the *z*-statistics are assumed to be independent in our simulations. We recommended pruning of SNPs (such that SNPs are roughly in linkage equilibrium) before applying our method of heritability estimation, however residual correlations may still exist. How the residual correlations may affect the SE estimates remains an open question.

The above resampling methods can potentially be speeded up by splitting the job into multiple processes to be run in parallel, although this approach has not be implemented in our software yet. We have not yet fully evaluated the building of confidence interval (CI) in our study but a natural approach is to assume normality and calculate CI in the form of 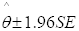. Assuming a polygenic model, the total heritability is the sum of Vg contributed by many variants of small to modest effect sizes. It is hence reasonable to assume normality by the central limit theorem. Further research may focus on developing other methods of building CIs and their comparisons.

## ACKNOWLEDGEMENTS

### Funding

This work was supported by the Hong Kong Research Grants Council General Research Fund grant and the University of Hong Kong Strategic Research Theme of Genomics.

